# Comprehensive study of *Trypanosoma cruzi* genetic diversity from Triatominae vectors in the Southern United States: Geographic structuring, mitochondrial introgression, and multiclonality

**DOI:** 10.64898/2026.08.21.746190

**Authors:** Juan C. Hernandez-Valencia, Norman L. Beatty, Kevin J. Vogel, Jan Zima, Eva Nováková

## Abstract

**Background:** *Trypanosoma cruzi*, the causative agent of Chagas disease, is subdivided into distinct genetic groups known as Discrete Typing Units (DTUs), each with distinct genetic traits that influence epidemiology and transmission dynamics. Several triatomine species serve as potential vectors of *T. cruzi* in the United States. However, despite the growing number of Chagas disease cases in the country, little is known about the genetic diversity and population structure of *T. cruzi* in natural vector populations.

**Methodology/Principal Findings:** We applied a multilocus metabarcoding approach to improve DTU resolution and characterize the genetic diversity and structure of *T. cruzi* in triatomines collected across five states of the southern United States. Five single-copy nuclear markers and one mitochondrial marker were amplified and processed by high-throughput sequencing to assess genetic diversity. We recovered 35 nuclear and 15 mitochondrial haplotypes from 70 infected specimens. Overall, genetic diversity was low (π < 0.01 at all nuclear loci), with DTUs TcI and the North American lineage of TcIV detected, TcI being the most prevalent. Geographic structuring was particularly evident in TcI strains, which exhibited a distinctive haplotype profile in Florida populations, potentially linked to the recently revalidated vector species *Triatoma ambigua*. Mitochondrial introgression from TcIV into TcI suggests inter-DTU genetic exchange in these populations. Multiple haplotypes within individual insects detected across single-copy nuclear markers, support multiclonal infection as common feature of *T. cruzi* in natural vectors.

**Conclusions/Significance:** These findings provide new insights into the genetic landscape and evolution of *T. cruzi* in the United States. Evolutionary connectivity through mitochondrial introgression and frequent multiclonality highlights the importance of deep sequencing approaches for resolving *T. cruzi* genetic diversity, with direct implications for understanding for transmission dynamics, disease monitoring and control.

**Author summary:** Kissing bugs are blood-feeding insects best known for transmitting *Trypanosoma cruzi*, the parasite that causes Chagas disease. The disease is usually associated with Latin America, and the United States is not recognized as an endemic country. Nonetheless, the insects live across the southern states, and people there have become infected without traveling abroad. Surprisingly little is known about the parasites themselves in this region. We collected kissing bugs from five southern states and sequenced six genetic markers using high-throughput sequencing to study the parasite variants present in those populations. We found parasites from Florida formed a genetically distinct group, and this separation aligns with a kissing bug species recently recognized as distinct. We also found that individual insects often carried several different parasite variants at once, and that some parasites combined genetic material from separate parasite groups. Characterizing *T. cruzi* infections improves our understanding of the ecological and epidemiological dynamics within hosts in the United States.

## Introduction

Chagas disease, caused by the protozoan parasite *Trypanosoma cruzi*, is a major vector-borne disease that affects millions of people in Latin America [1]. Traditionally linked to rural areas and poor housing conditions [2], its presence in the United States (U.S.) has been overlooked [3, 4], and only in recent years has it begun to receive growing attention from the scientific and medical communities [4–6]. The parasite is primarily transmitted by hematophagous insects in the subfamily Triatominae known most commonly as kissing bugs. While vector-borne transmission is well-documented in endemic regions of Latin America, growing evidence shows endemicity in the U.S. within certain regions where infected triatomine species are widespread, mostly in southern states such as Texas, California, and Gulf Coast regions [4, 6–9]. One model has estimated that about 288,000 people are currently infected in the U.S. who acquired infection while living or visiting Latin America, with around 10,000 cases attributed to autochthonous transmission within the U.S. [10]. Despite these findings, a tremendous gap in our understanding of the genetic diversity and evolutionary dynamics of *T. cruzi* in North America remains [11], especially compared with the extensive research conducted in Central and South America [12–15].

*Trypanosoma cruzi* exhibits substantial genetic diversity across its geographic range. The current classification framework recognizes this diversity as a series of genetically distinct Discrete Typing Units (DTUs). At least seven DTUs have been described (TcI-TcVI, and TcBat), and these lineages may differ in pathogenicity, host associations, and transmission dynamics [16–19]. In the U.S., TcI and TcIV are the predominant DTUs in mammalian reservoirs and triatomine vectors [11, 20]. TcI, the most widely distributed lineage in the Americas, is commonly associated with both domestic and sylvatic transmission cycles. In contrast, TcIV is primarily found in sylvatic cycles [18, 19]. Notably, TcIV in North America (designated as TcIV-North or TcIV-USA) shows a significant genetic divergence from its South American counterpart, suggesting it has followed an independent evolutionary path [20]. This divergence implies that TcIV-North has been evolving separately for over 100,000 years [20]. Despite this accumulated knowledge, further research is needed to explore the population structure and genetic diversity of the DTUs circulating among Triatominae populations in the U.S.

Recent studies have highlighted the genetic exchange and mitochondrial introgression between TcI and TcIV-North in the U.S [21–23]. Phylogenetic analyses have identified genetic recombination, with nuclear and mitochondrial markers showing conflicting DTU classifications. For example, isolates from Triatominae species have shown mitochondrial haplotypes clustering with TcI, while nuclear markers place them within TcIV-North, suggesting hybridization events [20, 23, 24]. These findings challenge the traditionally assumed clonal evolution of *T. cruzi* and highlight the need for broader genetic analysis.

Traditionally, most genetic studies on *T. cruzi* have relied on mitochondrial markers and multilocus sequence typing (MLST) performed through standard Sanger sequencing, which often overlook mixed infections, rare haplotypes, and recombinant lineages [25]. Recently, next-generation sequencing approaches have become more common, mainly targeting the mini-exon gene, a trypanosomatid-specific multicopy marker, widely used for detecting *T. cruzi* infection in epidemiological and diagnostic applications [26]. However, while multicopy genes are valuable for infection detection, the use of single- copy nuclear genes, which enables the analysis of orthologous sequences, is more suitable for phylogenetic purposes and genetic diversity studies of the parasite, allowing reproducible results [27–29]. Given the growing evidence of the complex genetic characteristics of *T. cruzi*, analyzing high-resolution single-copy nuclear markers using deep sequencing is key to refining our understanding of *T. cruzi* diversity in North America.

In this study, we apply a multilocus metabarcoding approach using deep sequencing to explore the genetic diversity and population structure of *T. cruzi* in triatomine species from the southern U.S. Specifically, we introduce three novel nuclear single-copy markers along with two well-established markers to enhance phylogenetic resolution and improve DTU genotyping, as well as a mitochondrial marker for detecting introgression. By combining nuclear and mitochondrial markers, this study aims to: i) Examine the genetic diversity and structure of *T. cruzi* populations in U.S. Triatominae vectors. ii) Identify signs of mitochondrial introgression and multiclonality within these populations, and iii) compare the effectiveness of Sanger sequencing versus NGS in characterizing *T. cruzi* genetic diversity. Using high-throughput sequencing technologies and a refined MLST framework, this work offers new insights into the complex genetic landscape of *T. cruzi* in the U.S.

## Methodology

### Sample collection and *T. cruzi* screening

Triatomine species were collected from different localities across the southern U.S. over a six-year period from 2017 to 2023. In Arizona, California, and Texas, destructive sampling of host nests was performed, while in Georgia and Florida, triatomines were collected from woodpiles and chicken coops in peridomestic environments where their presence was previously recorded. All individuals were preserved in absolute ethanol prior further processing. The DNA was extracted from the abdomen using the DNeasy Blood & Tissue Kit (QIAGEN) according to the manufacturer’s protocol. Species identity based on morphological characteristics was molecularly confirmed by cytB sequencing as published previously [30]. Each sample was screened for the presence of *T. cruzi* using PCR with primers TCZ1 and TCZ2, which target the mini-exon region [31]. The PCR products were visualized using gel electrophoresis, and samples displaying the expected band were considered positive for *T. cruzi*.

### Marker selection, PCR protocol and Library preparation

This study employed six genetic markers: one mitochondrial marker (cytb) and five nuclear markers. Two nuclear markers, *pumilio RNA-binding protein 1* (TcPUF1) and *pyruvate dehydrogenase kinase* (TcPDK), have been described previously [27], whereas three novel nuclear markers, designated 17c, 18c, and 17d, were developed in this study. To identify suitable loci, single-copy orthologous genes were screened across publicly available *T. cruzi* genomes in TriTrypDB release 59 (https://tritrypdb.org). Orthologous sequences were retrieved, supplemented with homologous sequences from the outgroups *T. rangeli* and *T. cruzi* marinkellei, and aligned in Geneious Prime v2023.2 [32] using MUSCLE plugin. Preliminary maximum-likelihood phylogenies were reconstructed to identify loci with sufficient sequence variation to discriminate among DTUs. Primers were subsequently designed in conserved regions flanking informative sites to maximize amplification across *T. cruzi* diversity while minimizing cross-amplification of related trypanosomes (Table 1). Primer performance was validated experimentally using *T. cruzi*-positive samples and *T. rangeli* as a specificity control. The best-performing markers, 17c, 18c, and 17d, target single-copy hypothetical genes in the *T. cruzi* YC6 reference genome, corresponding to locus tags TcYC6_0032630, TcYC6_0017400, and TcYC6_0030750, respectively.

**Table 1.**
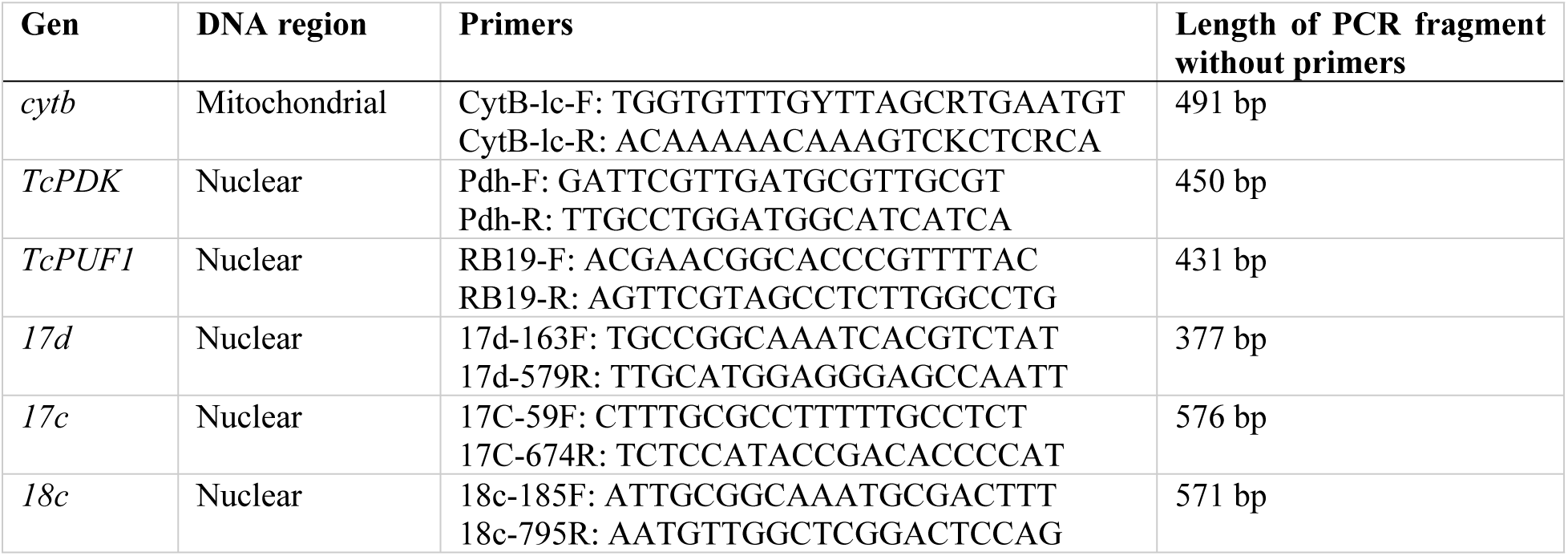
Information of the six genes and corresponding primers used in this study.

Each marker was individually amplified using the QIAGEN Multiplex PCR Kit. An initial amplification was performed separately for each marker, and the amplified products were visualized on an agarose gel to confirm successful amplification. Each reaction consisted of 12 µl Master mix, 1 µl of each primer, 1.5 µl of DNA and 8.5 µl of PCR water. PCR conditions: 15 min of initial denaturation followed by 35 cycles of denaturation (94°C) for 30 s, annealing (50°C for *17d* and *cytb*, 55°C for *TcPDK*, *TcPUF1*, *17c* and *18c*) for 90 s, elongation at 72°C for 45 s, with a final elongation step for 10 min. The six PCR products were pooled and used as templates in a second PCR to incorporate sample-specific barcodes and Illumina adapters. Each 40-µL reaction contained 20 µL of Q5 PCR Master Mix, 4 µL each of barcoded forward and reverse primers (2.5 nM), 2 µL of the first-round PCR product, and 10 µL of Milli-Q water. Amplification comprised an initial denaturation at 94°C for 3 min, followed by 12 cycles of denaturation at 94°C for 45 s, annealing at 65°C for 20 s, and extension at 72°C for 30 s, with a final extension at 72°C for 2 min. The barcoded PCR products were visualized on a gel, purified using SPRIselect beads (Beckman Coulter), and quantified using a spectrophotometric microplate reader Synergy H1 (BioTek). The quantified products were then pooled in equimolar amounts to ensure uniform representation in subsequent analyses.

The control AMPLIRUN® *Trypanosoma* DNA Control (MBC059-R), derived from the *T. cruzi* Y strain, was processed along with the samples to ensure the accuracy and reliability of the sequencing methods.

### High-Throughput Sequencing and data analysis

Purified amplicon library for all six markers was sequenced on an Illumina MiSeq platform (MiSeq Reagent Nano Kit v2, 250 bp paired-end reads). Demultiplexed sequencing data were processed in USEARCH v11 [33]. Paired-end reads for *17d*, *TcPUF1*, and *TcPDK* were merged using fastq_mergepairs (minimum overlap 16 bp). For *17c*, *18c*, and cytb, whose amplicons do not reach the minimum overlap length, reads were joined using fastq_join, which reverse-complements the reverse read and inserts padding between reads. A minimum merged/joined length of 300 bp was required in both cases. Forward primers were used to extract sequences corresponding to each marker. Subsequently, primers were removed, and the sequences were quality-filtered and truncated to the expected lengths. To remove chimeric sequences and low-quality assemblies, zero-radius operational taxonomic units (ZOTUs) were defined using the UNOISE algorithm [34]. Each ZOTU was screened against *T. cruzi* genomes available in the NCBI database using BLAST queries [35]. To further reduce sequencing artifacts, including errors caused by cross-talk [33], only sequences representing at least 0.25% of the total reads in each sample were retained. Other thresholds of 0.1% and 0.5% were tested, with no major differences in the output sequences.

### Phylogenetic analyses

The sequences used for phylogenetic analyses included ZOTUs generated in this study and reference sequences retrieved from *T. cruzi* whole-genome assemblies available in NCBI [36] and TriTrypDB [37] (S1 Table). Additionally, raw whole genome sequencing data from the *T. cruzi* CanIII strain (BioProject accession: PRJNA198812), which did not have a preassembled genome available in those repositories, were mapped to the selected marker sequences identified in the other available genomes. The data were subsequently assembled for inclusion in phylogenetic analyses. Sequences from GenBank were also incorporated into the dataset for previously characterized markers *TcPUF1*, *TcPDK*, and *cytb*.

For each of the six genetic markers, sequences were aligned using MUSCLE v.3.8.425 [38] with the default settings. The resulting alignments were manually checked to avoid misalignments. The best-fit substitution model for each alignment was determined using IQ-TREE’s ModelFinder with the Bayesian Information Criterion (BIC) to select the most appropriate model for each dataset. Phylogenetic trees were inferred using IQ-TREE v.2.2.0 [39]. The chosen substitution model for each marker was used in the maximum-likelihood (ML) analysis. To assess the robustness of the inferred phylogenies, bootstrap analysis was conducted with 1000 replicates. In addition, a Bayesian phylogenetic inference was performed using MrBayes v.3.2.7 [40]. The selected substitution model was applied, and Markov Chain Monte Carlo (MCMC) analysis was carried out with four chains for 1000000 generations, sampling every 1000 generations. The standard deviation of split frequencies was monitored to ensure convergence, with a burn-in period set to 25% of the total generations.

To construct a phylogeny from concatenated data across all nuclear markers, we first obtained a consensus sequence for each marker by locality, species, and DTU. This approach produced a single representative sequence for concatenation. Since the haplotypes were not highly divergent and consistently clustered within the same branches in the individual marker trees, combining them into consensus sequences is biologically justified. These consensus sequences accurately reflect the underlying phylogenetic signal, and the minor sequence variations do not significantly affect the overall phylogenetic relationships. The consensus sequences were then concatenated and aligned with sequences derived from publicly available *T. cruzi* genomes using Geneious v.2022.0.1. The concatenated alignment underwent the same model selection and phylogenetic analysis as those used for the individual markers described above. For both the individual marker trees and the concatenated tree, sequences from the *T. cruzi* marinkellei strain were designated as the outgroup. This outgroup was selected based on prior phylogenetic knowledge and was consistently applied across all analyses [28].

### Genetic diversity and structure analysis

Haplotype networks were constructed for each marker using the Templeton, Crandall, and Sing (TCS) method using PopART v. 1.7 [41]. To evaluate mutation-drift equilibrium and assess genetic diversity, we performed Tajima’s D neutrality test, and estimated nucleotide diversity (π), polymorphic sites, and parsimoniously informative sites using PopART and DNAsp v. 6 [42].

Genetic differentiation among samples was visualized using principal component analysis (PCA) based on a genetic distance matrix and statistically assessed with permutational multivariate analysis of variance (PERMANOVA) with 10000 permutations. This analysis aimed to determine whether genetic structure was significantly associated with factors such as the geography or triatomine species. Additionally, we conducted an analysis of molecular variance (AMOVA) to further explore the influence of these factors on genetic structure. To test population structure across various groups, we evaluated the significance of the Φst and Fst indices of genetic differentiation using permutation tests with 10000 replicates. All statistical analyses, including PCA, PERMANOVA, and AMOVA, were performed using the stats and vegan packages in R [43, 44]. A Benjamini-Hochberg false discovery rate (FDR) correction was applied to p-values from AMOVA and Φst analyses, with significance maintained at FDR < 0.05.

### Comparison of Sanger and Metabarcoding sequencing

Sanger sequencing was used to compare the effectiveness of the two approaches (multilocus metabarcoding and Sanger sequencing) for characterizing the genetic diversity of *T. cruzi* in vectors. This analysis aimed to identify discrepancies between the methods and evaluate their resolution in detecting genetic variation. The *17d* marker was sequenced in 45 specimens, while the *17c* and *18c* markers were sequenced in 18 specimens each. Additionally, eight specimens were processed for the *TcPDK* marker, and ten specimens for the *TcPUF1* marker, while the mitochondrial marker *cytb* was sequenced for six specimens. All sequences generated through Sanger sequencing were integrated with those obtained via metabarcoding to construct phylogenetic trees and compare genetic diversity metrics.

The same workflows used for metabarcoding-derived sequences were also applied to Sanger-generated sequences, with a focus on the *17d* marker to facilitate direct comparisons. Sanger sequencing was conducted at SEQme (Czech Republic) using the same primers and protocols previously described for high-throughput sequencing, thereby ensuring consistency across methods.

## Results

Over 6 years of sampling, 1,130 individuals were screened for *T. cruzi*. Among these, 495 individuals (43.8%) tested positive for the parasite. For further genetic analysis, 72 individuals collected in California (n = 3), Arizona (n = 22), Texas (n = 26), Georgia (n = 4), and Florida (n = 17) were selected (S2 Table), representing five species of triatomines: *Triatoma gerstaeckeri*, *T. rubida*, *T. sanguisuga* s.l., *Hospesneotomae protracta,* and *Paratriatoma lecticularia* (Fig 1A). We retrieved amplicon sequencing data for 70 individuals, comprising 621,733 reads, with an average of 9,011 ± 3,611 reads per sample.

**Fig 1.**
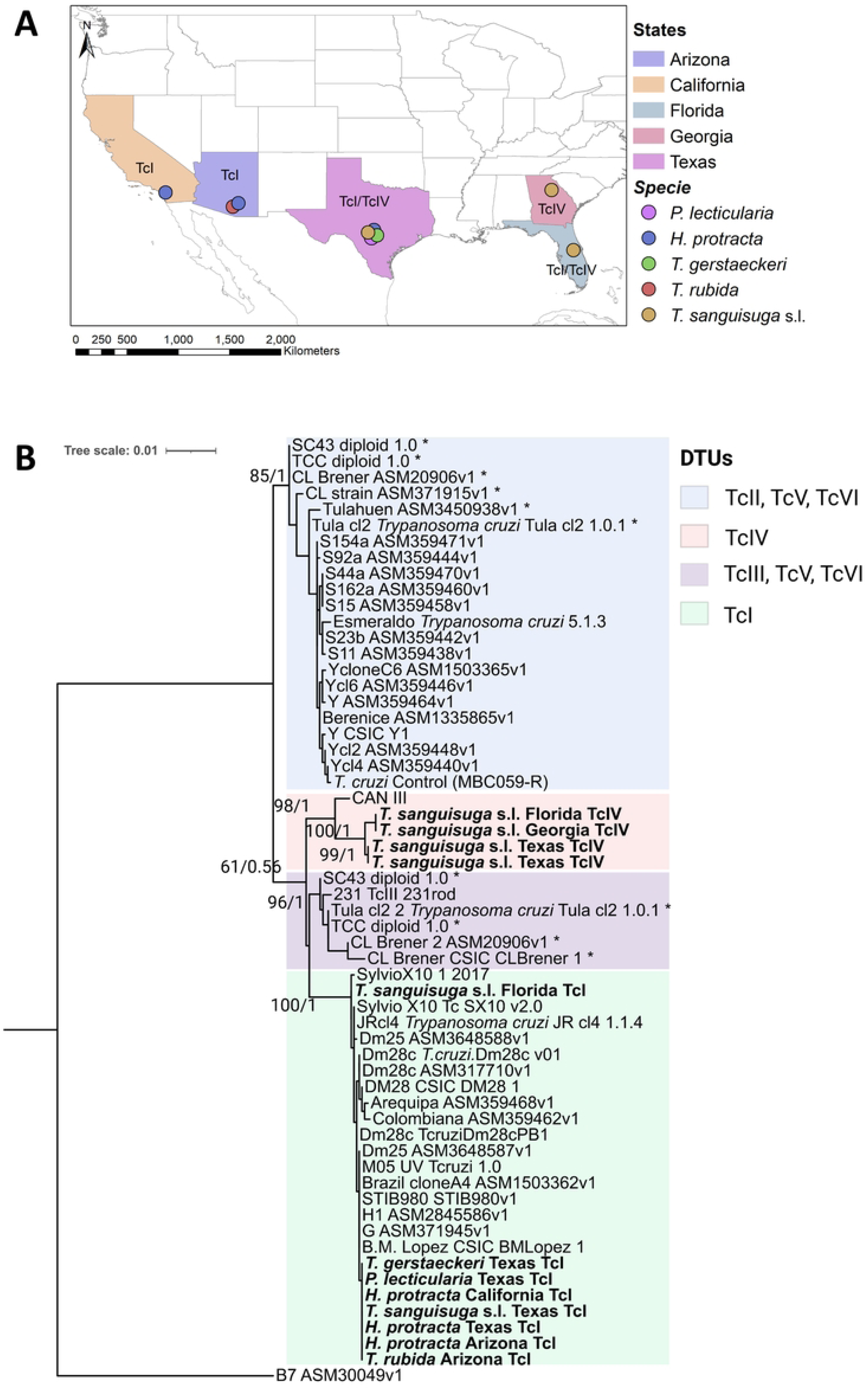
Sampling locations of Triatominae species and *T. cruzi* phylogeny based on nuclear loci concatenated data. (A) Map showing the distribution of triatomine species and their collection sites across the southern U.S. (B) Phylogeny of *T. cruzi* derived from concatenated sequences of five nuclear markers. Consensus sequences for each marker were grouped by state, species, and identified DTU; sequences followed by an asterisk (*) correspond to assemblies from hybrid DTU strains. Branch values indicate the statistical support from the maximum likelihood analysis (first value) and the posterior probability from the Bayesian inference (second value).

### Phylogenetic analysis of nuclear markers

Phylogenetic trees generated from individual nuclear markers consistently recovered the recognized *T. cruzi* DTUs (S1–S5 Figures). The three novel markers (*17c*, *17d*, and *18c*) provided strong phylogenetic resolution, clearly distinguishing TcI, TcII, TcIII, and TcIV, and performed comparably to already established marker *TcPUF1*. In contrast, the previously published *TcPDK* marker was more conserved and showed lower discriminatory power, resulting in polytomy among the main DTUs (S4 Figure). Except for *TcPDK*, all markers supported monophyly of TcI–TcIII–TcIV cluster as a sister clade to TcII, demonstrating the suitability of the newly developed markers for DTU discrimination. The hybrid DTUs TcV and TcVI occupied their expected positions within the parental TcII and TcIII lineages.

The final concatenated dataset for the five nuclear markers comprised 70 samples from this study and 46 reference sequences derived from publicly available *T. cruzi* genomes (S1 Table), yielding an alignment of 1,910 bp. The resulting phylogeny was strongly supported across all major nodes (bootstrap >90%, posterior probability >0.99; Fig. 1B). Consistent with the individual-marker analyses, TcII formed a distinct lineage sister to a major clade comprising TcIV, TcIII, and TcI. Within this clade, TcI and TcIII were recovered as sister lineages, with TcIV branching basally to them.

All *T. cruzi* lineages detected in U.S. triatomines were assigned to either TcI or TcIV-North, confirming the DTU assignments obtained from individual-marker analyses. *T. cruzi* TcI from vectors collected in California, Arizona, and Texas formed closely related lineages, whereas the Florida TcI sequences occupied a different position within the TcI clade. TcIV-North haplotypes clustered with the reference CANIII strain in a well-supported clade and showed evidence of internal substructure possibly corresponding to geographic origin. The Y-strain positive control consistently clustered within the TcII clade (Fig. 1B).

### Phylogenetic analysis of mitochondrial marker (*cytb*)

Aligning *T. cruzi* mitochondrial cytb fragments retrieved in this study, along with reference sequences from GenBank, resulted in a matrix of 378 nucleotide positions for a total of 279 sequences. Phylogenetic analysis of these sequences recovered five well-supported mitochondrial lineages (Fig 2A). One major clade grouped sequences associated with the TcIII and TcIV-South DTUs, as well as the hybrid DTUs TcV and TcVI. This clade was a sister group to a distinct TcIV-North lineage. The remaining lineages corresponded to TcI, TcBat, and TcII, each forming well-defined and separate clades. All *T. cruzi cytb* sequences obtained from natural populations of triatomines in the U.S clustered within either the TcI or TcIV-North lineages. Within TcI, nine mitochondrial haplotypes were identified, whereas six haplotypes were detected within TcIV-North. The TcIV-North haplotypes grouped together with the CANIII reference strain and remained clearly separated from the South American TcIV lineage. Consistent with the nuclear- marker analyses, no U.S. samples were assigned to TcII, TcIII, TcV, or TcVI.

**Fig 2.**
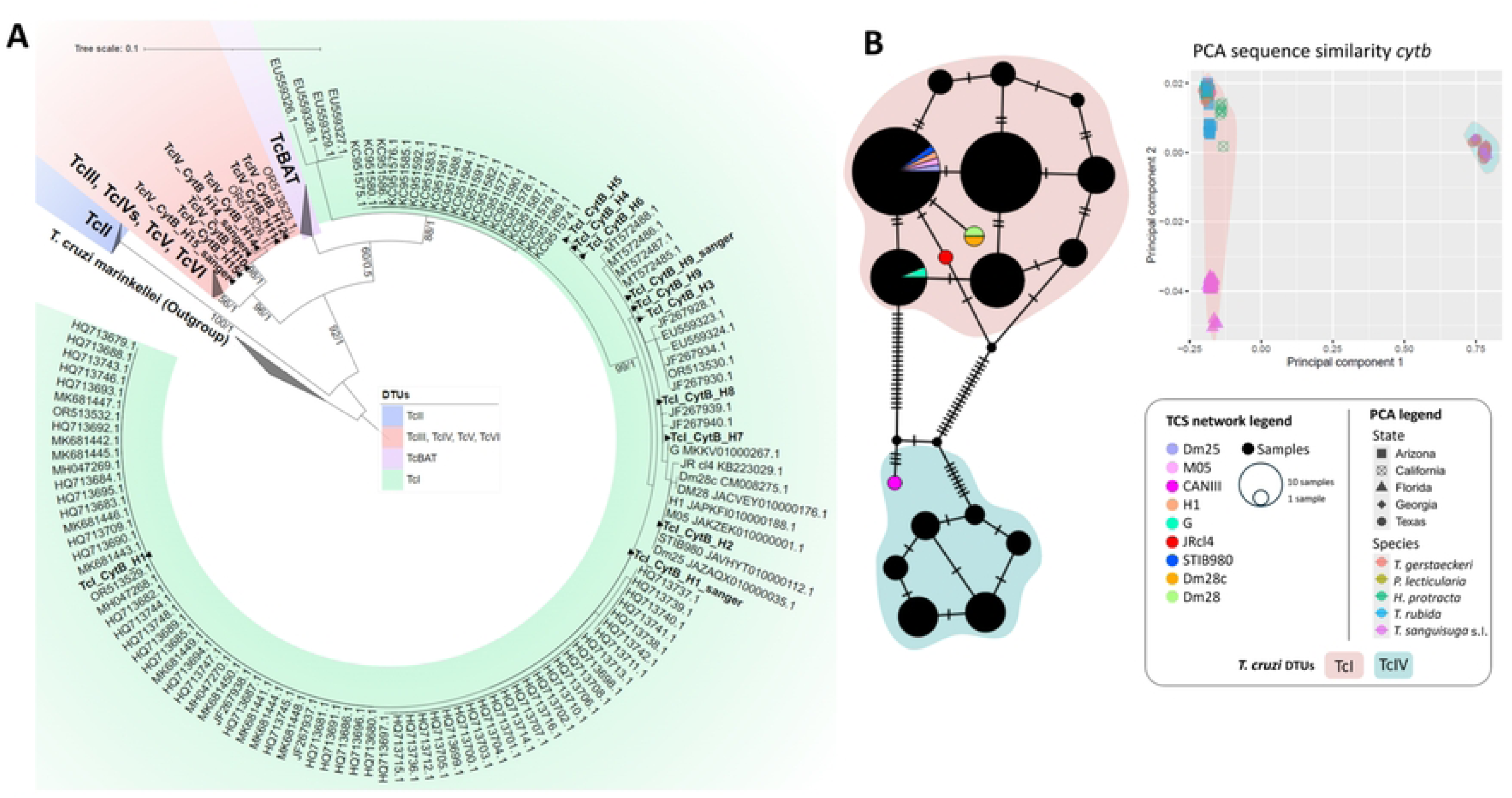
Phylogeny based on cytochrome B (*cytb*) sequences, haplotype network, and PCA plot for *T. cruzi* in Triatominae species from the southern U.S. (A) Phylogeny of *T. cruzi* derived from *cytb* sequences. Branch values represent statistical support from the maximum likelihood analysis (first value) and the posterior probability from Bayesian inference (second value). The *cytb* sequences identified in this study are highlighted in bold within the tree. (B) Haplotype TCS networks and PCA plots based on sequence similarity of *cytb*. Haplotypes identified in this study are represented in black, while reference sequences extracted from *T. cruzi* whole-genome sequencing (WGS) data are shown in color.

### Phylogenetic insights and haplotype relationships

The TCS haplotype network and phylogenetic analysis provided valuable insights into the genotypic composition of *T. cruzi* detected in this study. All haplotypes identified as DTU TcI across the nuclear loci grouped with references belonging to the TcId genotype. In contrast, DTU TcIV exhibited genetic divergence from the reference strain CANIII, which represents the TcIV-South genotype. This divergence was evident in both the phylogenetic analysis and the TCS haplotype network (Figs 1 and 3). However, an exception was identified for locus *TcPUF1*, where one haplotype (TcIV_RB_H4) found in *T. sanguisuga* s.l. from Georgia was identical to the CANIII reference (Fig 3E).

**Fig 3.**
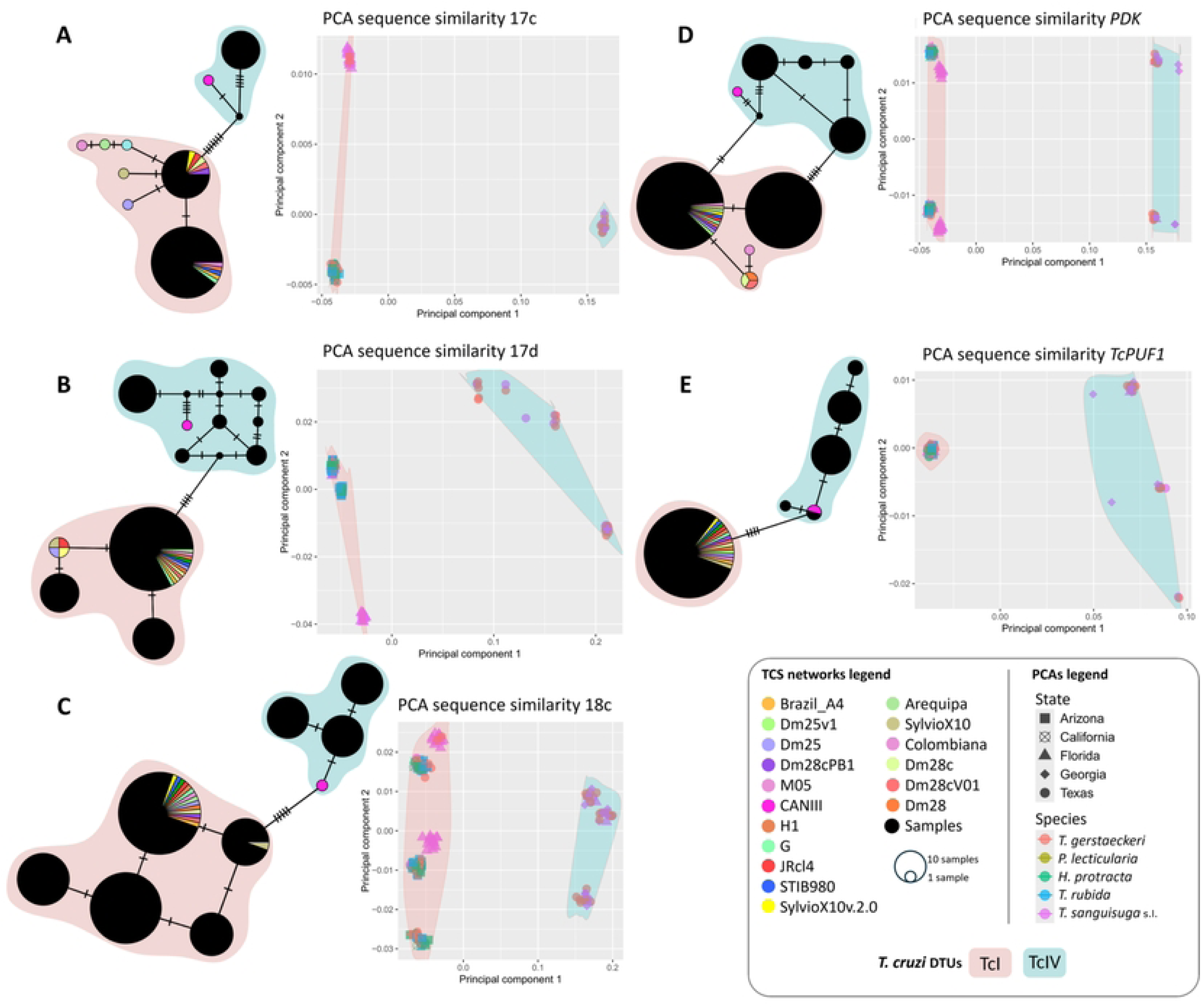
Haplotype networks and PCA plots based on sequence similarity for the five nuclear markers of *T. cruzi* in Triatominae species from the southern U.S. The figure shows haplotype networks constructed using the TCS method and PCA plots based on sequence similarity for each of the five nuclear markers analyzed. The haplotypes identified in this study are represented in black, while reference sequences extracted from *T. cruzi* whole-genome sequencing data are shown in color. (A) *17c*, (B) *17d*, (C) *18c*, (D) *TcPDK*, and (E) *TcPUF1*. The TCS networks shown here provide context for our haplotypes within known *T. cruzi* DTUs. The sequence trimming required for alignment with database references resulted in the loss of some nucleotide positions, reducing resolution and causing certain study haplotypes to merge. The complete, untrimmed networks retaining full resolution are shown in Figure S6.

The phylogenetic analysis corroborated these findings. Specifically, loci *17c*, *18c*, *17d*, and *TcPDK* showed distinct clades for the TcIV-North and TcIV-South genotypes, while locus *TcPUF1* displayed one haplotype clustering with the CANIII reference (Fig 3E and S5 Fig). Conversely, the concatenated phylogeny, constructed using consensus sequences from each species across different states, revealed a clear separation between the TcIV-South and TcIV-North genotypes (Fig. 1B).

Additionally, it is noteworthy that the combined analysis of nuclear loci and the mitochondrial marker indicated mixed infections involving TcI and TcIV in three specimens. Furthermore, three specimens from Texas, corresponding to the species *T. gerstaeckeri* and *T. sanguisuga* s.l., exhibited phylogenetic incongruence between the nuclear loci and the mitochondrial marker. While the nuclear loci indicated only TcI, the *cytb* marker revealed a mitochondrial profile associated with TcIV.

### Genetic diversity and structure

Across the samples, a total of 35 unique genetic sequences were identified from all analyzed nuclear loci (S3 Table). Among these loci, *17c* showed the lowest haplotype number, with three haplotypes. This was followed by loci *17d* and *TcPDK*, with ten and eight haplotypes, respectively. Conversely, locus 18c displayed the highest haplotype diversity, with eight haplotypes identified (Table 2). It is noteworthy that loci *17d* and *17c* consistently showed no more than two different haplotypes per sample. In contrast, other loci exhibited more than two haplotypes. Specifically, locus *18c* showed more than two haplotypes in 33 samples, *TcPDK* in 13, and *TcPUF1* in 3.

**Table 2.**
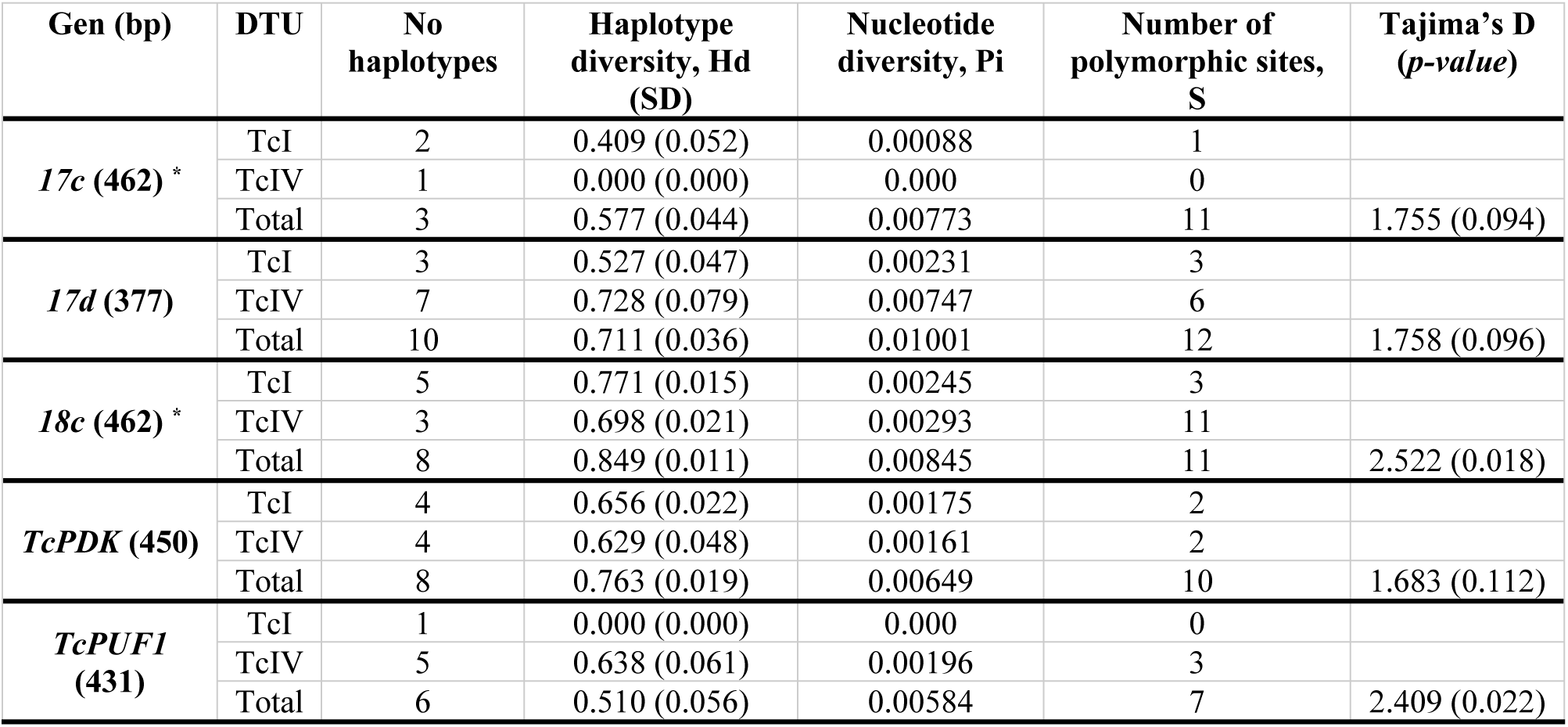
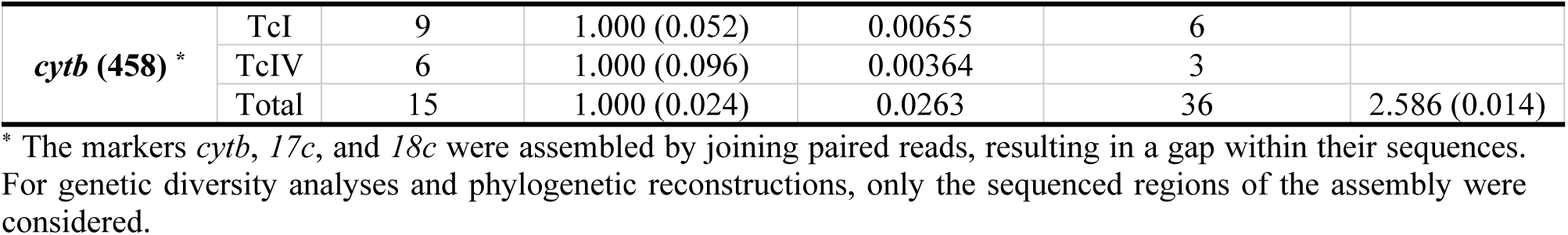
Genetic diversity indices for *T. cruzi* from Triatominae species across the southern U.S.

The number of unique haplotypes varied among the different loci, but the analyses of genetic diversity and structure revealed consistent patterns. The overall nucleotide diversity across the dataset indicated low genetic variation (π < 0.01). This was especially evident at the intra-DTU level, where genetic diversity was markedly lower. Tajima’s D neutrality tests consistently yielded positive values across all loci (D > 1.68), indicating an excess of common genetic variants in samples from different states and species (Table 2). While not statistically significant across all loci, this pattern is consistent with the predominant clonal nature of *T. cruzi*, which tends to preserve shared genetic variants within DTUs.

Haplotype networks constructed using TCS for each nuclear locus displayed two distinct haplotype groups, which were separated by varying numbers of mutational steps: four for *TcPUF1* and *17d*, seven for *18c* and *TcPDK*, and ten for *17c* (S6 Fig). These groups correspond to the DTUs TcI and TcIV. Results from AMOVA confirmed that the DTU was the primary factor explaining genetic variation among *T. cruzi* populations within triatomine species across the southern U.S. (R² = 0.87-0.99, *p* < 0.001). However, while DTU differences accounted for most of the genetic variation, geographic patterns of haplotype differentiation were evident at the intra-DTU level (S5 Table).

For TcI, distinct haplotypes were consistently associated with specific states across most loci. Unique haplotypes were identified in Florida for all loci except *TcPUF1*, while an exclusive haplotype for locus *17d* was observed in Arizona (S6 Fig). These findings were supported by AMOVA and Φ-statistics analyses, which indicated that geographic origin significantly explained part of the genetic variance in TcI (R² = 0.14-0.77, *p* < 0.001; Φst = 0.41-0.55, *p* < 0.001), demonstrating moderate but statistically significant geographic differentiation, except for the *TcPUF1* locus (*p* = 0.14). In contrast, for TcIV, unique haplotypes were identified in Georgia (*TcPDK* and *TcPUF1*) and Texas (*TcPUF1* and *17d*), but no statistical evidence of geographic differentiation was observed (S6 Fig and S5 Table). This suggests a greater genetic homogeneity among *T. cruzi* TcIV populations in Texas, Georgia, and Arizona, where this DTU was found.

In addition to geographic origin, the species factor contributed moderately to the observed genetic variation (R² = 0.05-0.55, *p* < 0.001; Φst = 0.10-0.38, *p* < 0.05) (S5 Table). While some degree of genetic structuring was evident among Triatominae species, the TCS networks showed haplotypes that were exclusive to *T. sanguisuga* s.l., whereas other haplotypes were shared among multiple species (S6 Fig). This pattern likely reflects the geographic distribution of *T. sanguisuga* s.l., which was the only species collected in Florida, a state with a distinct haplotype profile in most loci (S6 Fig and S3 Table).

Principal component analysis provided a clear visualization of genetic differentiation among samples for all loci, with the first two principal components explaining over 99% of the variance. The observed clustering, corroborated by PERMANOVA results, supported the genetic structure identified through AMOVA and TCS network analyses (Fig 3). Both DTU and geographic origin emerged as dominant factors shaping the genetic structure of *T. cruzi* within Triatominae natural populations in the southern U.S.

The *cytb* mitochondrial marker, analyzed across 458 nucleotide positions in 70 samples, showed higher nucleotide diversity than the nuclear loci, with π = 0.026, indicating a moderate level of genetic variation at the mitochondrial level. A total of 36 segregating sites were identified, of which 35 were parsimony informative. The genetic structure observed for this mitochondrial locus was similar to that of the nuclear loci. The Tajima’s D test for *cytb* yielded a positive value (D = 2.59, *p* < 0.05), suggesting an excess of common haplotypes across triatomine species from the southern U.S. (Table 2). The TCS haplotype network displayed two distinct haplotype groups corresponding to the DTUs TcIV and TcI, separated by 29 mutational steps (Fig 2). AMOVA and PERMANOVA confirmed that DTU was the primary factor explaining genetic variation among *T. cruzi* populations within triatomine species across the southern U.S. (R² = 0.98, *p* < 0.001). However, at the intra-DTU level, geographic origin was the main factor explaining genetic variation for TcI (R² = 0.82, *p* < 0.001), with specific haplotypes observed in California, Arizona, and Florida. For DTU TcIV, no genetic structuring was observed among specimens from the states where it was detected (R² = 0.08, *p* > 0.05) (S5 Table).

### Comparison of Sanger and Metabarcoding sequencing

Specimens processed through Sanger sequencing showed differences in the number of haplotypes compared to the metabarcoding method for each marker. Sanger sequencing consistently identified the dominant haplotype from each specimen, while metabarcoding detected multiple haplotypes per marker within the same sample (S3 and S4 Tables). In some cases, Sanger produced ambiguous signals at variable positions among co-occurring haplotypes. This ambiguity was particularly evident for the *17d* marker in specimens collected in Florida. While metabarcoding identified two distinct haplotypes for this marker, Sanger sequencing yielded an ambiguous consensus sequence with undefined signals in positions where the haplotypes differed (data not shown). However, this ambiguity was not observed in all samples where metabarcoding detected multiple haplotypes.

As expected, direct Sanger sequencing failed to detect mixed infections involving distinct DTUs identified by metabarcoding. Specifically, metabarcoding retrieved haplotypes corresponding to both DTUs TcI and TcIV for each marker, whereas Sanger sequencing identified only a single haplotype per marker, all associated with DTU TcIV.

The genetic diversity analysis of the *17d* marker, which had the most extensive coverage among Sanger-sequenced markers, showed lower nucleotide diversity (π = 0.001) than the metabarcoding approach (π = 0.010). The Sanger dataset also displayed fewer segregating sites, and the Tajima’s D neutrality test returned a significantly higher positive value for the Sanger dataset (D =26.78), suggesting a greater excess of common genetic variants compared to metabarcoding. Additionally, the number of haplotypes was lower in the Sanger dataset. The TCS haplotype network based on Sanger sequences identified two distinct groups corresponding to DTUs TcI and TcIV, separated by only two mutational steps. In contrast, the haplotype network derived from metabarcoding revealed unique haplotypes, some of which were exclusive to specific states, reflecting greater genetic complexity, multiclonality, mixed infections, and localized variants (Fig 4).

**Fig 4.**
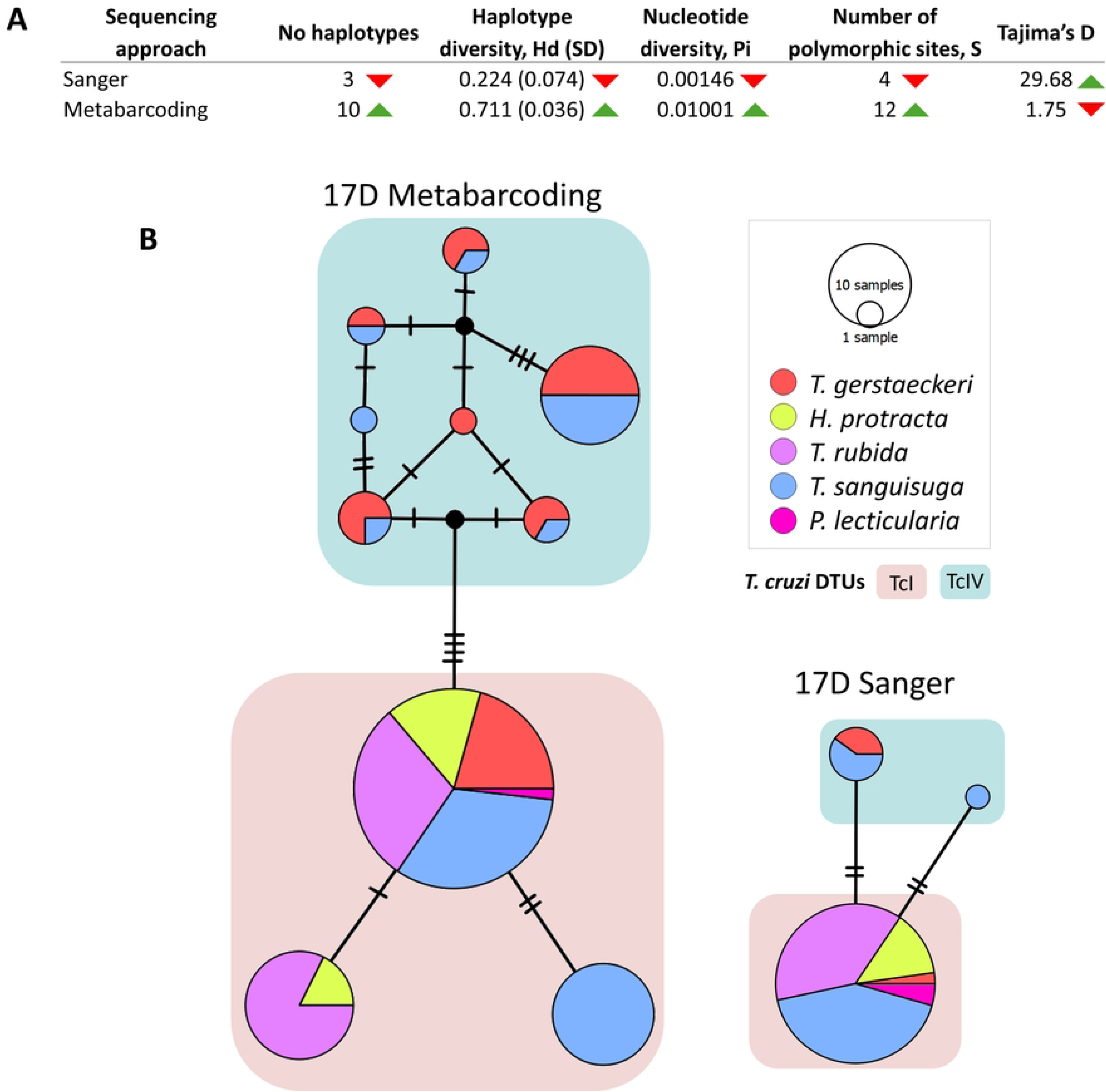
Comparison between Sanger and Metabarcoding sequencing for *17d* nuclear marker. (A) Genetic diversity indices for *17d* nuclear marker. (B) TCS haplotype networks based on metabarcoding (left) and Sanger (right) approaches.

## Discussion

Our study explores the genetic diversity of *T. cruzi* across different Triatominae populations in the southern U.S. over a six-year period. Applying a metabarcoding approach, we achieved a detailed characterization of the genetic complexity within these vector populations. Furthermore, we introduced three novel single-copy nuclear markers that demonstrate proper phylogenetic resolution, facilitating direct comparisons with reference strains available in public databases.

Phylogenetic analysis of these single-copy genes effectively resolved the distinct *T. cruzi* DTUs, revealing a phylogenetic structure that clearly distinguishes TcI, TcII, TcIII, and TcIV. As expected, DTUs TcV and TcVI grouped with TcII and TcIII, respectively, consistent with their hybrid origins [45]. These clustering patterns align with previously established findings [20, 21, 27, 46–48]. Our results reiterate these relationships and underscore the utility of the novel nuclear markers for improving genetic typing and evolutionary studies, and for refining assessments of *T. cruzi* diversity.

Phylogenetic typing using both nuclear and mitochondrial markers indicated that all specimens analyzed in this study belonged to DTUs TcI and TcIV. The haplotypes consistently clustered within clades corresponding to reference sequences from these DTUs, supporting accurate classification. Notably, TcIV exhibited clear geographic differentiation, with North American haplotypes forming a distinct cluster separate from South American haplotypes, consistent with previous findings [20, 48, 49]. The consistent identification of TcIV in our study, as a distinct and well-supported lineage across both nuclear and mitochondrial markers, supports previous proposals that TcIV-North constitutes an independently evolving lineage, rather than a geographic variant of South American TcIV [20]. The observed stability of this lineage across diverse hosts, vectors, and geographic regions suggests a prolonged evolutionary history within North American transmission cycles, likely exceeding the timescales anticipated by recent dispersal scenarios. These findings contribute to the growing body of evidence suggesting that the current discrete typing unit framework may underestimate the evolutionary distinctiveness of TcIV-North [20]. No other DTUs were detected in this study, a result consistent with previous research on U.S. triatomine populations, where DTUs other than TcI and TcIV-North have rarely been observed, reaffirming the predominance of these two lineages in the region [20].

The analysis of the studied genetic markers showed low genetic diversity among *T. cruzi* populations in the southern U.S., characterized by a predominance of shared genetic variants across different states and Triatominae species. This pattern reflects the predominantly clonal nature of *T. cruzi*, which maintains conserved genetic variants within distinct DTUs [50]. However, within the DTU TcI, geography significantly influenced the genetic variance. Unique haplotypes were identified in Texas, Georgia, and Arizona, although this geographic differentiation was particularly pronounced in Florida, where distinct haplotypes were detected across all loci except for *TcPUF1*. These results align with previously reported geographic patterns of *T. cruzi* genetic differentiation [51–53], emphasizing the role of biogeographical factors in shaping population structure.

The unique genetic profile of TcI populations in Florida, compared to those from Arizona, Texas, and California, may be attributed to several factors. A plausible explanation is the “peninsula effect,” where geographic isolation limits gene flow and influences demographic processes [54]. Florida’s limited land connection to the mainland could act as a bottleneck, restricting the introduction of new genetic variants and promoting genetic divergence [55]. This pattern of differentiation between peninsular and mainland populations has been observed in different species [56–58]. The distinct TcI haplotype detected in Florida may reflect not only geographic isolation of *T. cruzi* populations but also their association with a distinct vector species in this region. Recent integrative taxonomic analyses, based on concordant molecular and morphological evidence, support the revalidation of *Triatoma ambigua* as a species distinct from *T. sanguisuga* in Florida [59]. The correspondence between the divergent Florida TcI lineage and the geographic distribution of *T. ambigua* suggests that vector diversification may help structure *T. cruzi* populations in the Florida peninsula. Differences in vector ecology, host associations, and transmission networks could promote long-term parasite divergence. Although additional sampling across the southeastern U.S. is needed, these findings highlight the potential role of vector evolutionary history in shaping contemporary patterns of *T. cruzi* genetic diversity.

The multilocus metabarcoding approach allowed for the detection of multiple haplotypes per sample, with most Triatominae specimens harboring two or more haplotypes on at least one marker. Parasites often coexist within hosts in complex infrapopulations composed of multiple genetic variants [60]. Multiclonality in *T. cruzi* has been widely documented across different host types [61, 62], including triatomine species [63, 64], suggesting that multiclonal infections are more frequent than those involving a single genotype [63, 65]. The frequent detection of multiple haplotypes within individual triatomines suggests that multiclonal *T. cruzi* infections may be more common in U.S. transmission cycles than previously recognized. Because the markers employed in this study are single-copy nuclear loci, the recovery of more than two haplotypes from a single specimen cannot be explained by diploidy alone and instead indicates the presence of multiple parasite genotypes within the vector. Such multiclonal infections likely arise through repeated blood feeding on infected hosts or exposure to hosts harboring genetically diverse parasite populations [65]. The coexistence of multiple *T. cruzi* lineages within individual vectors may have important epidemiological and evolutionary consequences, increasing opportunities for genetic exchange and facilitating the maintenance of parasite diversity within local transmission networks [65, 66]. This suggests that co-infections could dominate natural transmission cycles, a reality that was previously masked by the technical limitations of traditional genotyping tools.

Our study also identified mitochondrial introgression in three specimens, as demonstrated by phylogenetic incongruence between nuclear and mitochondrial markers from the same samples. While nuclear markers placed these samples within TcI, their mitochondrial sequences clustered with TcIV. Mitochondrial introgression in *T. cruzi* was first reported by Machado and Ayala (2001), who observed discordant phylogenies in strains classified as TcI and TcIV (formerly TcI and TcIIa). Since then, different studies have reported TcI-TcIV mitochondrial introgression across a range of hosts in North and South America [21–23]. Specific reports in vectors include *T. sanguisuga* s.l. from Florida [23], *H. protracta* from California [24], and both *T. gerstaeckeri* and *P. lecticularia* from Texas [20]. Our findings expand on this evidence, identifying TcI-TcIV mitochondrial introgression in *T. gerstaeckeri* and *T. sanguisuga* s.l. from Texas. The detection of mitochondrial introgression indicates that, as in South American populations, contemporary *T. cruzi* diversity in the United States is shaped not only by divergence among DTUs but also by historical interactions among them [67, 68]. Rather than representing completely isolated evolutionary lineages, DTUs appear to retain the capacity for occasional genetic exchange, generating discordant mitochondrial and nuclear genealogies [69]. This evolutionary connectivity may be particularly important in regions where multiple DTUs co-circulate and mixed infections are common, creating opportunities for rare but consequential genetic exchange events. The maintenance of reproductive compatibility between DTUs, despite substantial genetic divergence, suggests that current DTU boundaries may be more porous than traditionally assumed and highlights the importance of considering *T. cruzi* as a dynamic, interconnected metapopulation rather than a collection of isolated lineages.

Finally, this study integrated metabarcoding and Sanger sequencing to compare their effectiveness in assessing *T. cruzi* genetic diversity. Our findings demonstrated that Sanger sequencing consistently recovered only the dominant haplotype from each specimen and, in some cases, generated ambiguous signals at polymorphic sites. Consequently, it failed to detect multiclonality and mixed infections. In contrast, deep sequencing provided a higher-resolution characterization of haplotypes per marker within the same specimen, yielding a more comprehensive view of *T. cruzi* infection. Comparative analyses of genetic diversity indices for the *17d* marker showed that Sanger sequencing overrepresented common genetic variants while underestimating overall genetic diversity. The ability of metabarcoding to detect rare haplotypes, multiclonality, and mixed infections offers a more accurate representation of *T. cruzi* genetic complexity, population structure, and evolutionary dynamics. This is particularly relevant from an epidemiological perspective, as genetic diversity potentially plays a key role in shaping parasite transmission dynamics [17, 25, 70]. Therefore, high-throughput sequencing technologies, whether applied to informative genetic markers or to genome-wide approaches, provide the most robust means of accurately characterizing *T. cruzi* genetic complexity in natural host populations, thereby enhancing our understanding of its epidemiological and evolutionary significance.

## Conclusion

Using a targeted metabarcoding approach to nuclear and mitochondrial markers, this study showed low genetic diversity of *T. cruzi* among Triatominae populations in the southern U.S., which were composed of DTUs TcIV and TcI, with the latter being the most widely distributed. Both DTU and geographic origin emerged as key factors shaping the genetic structure of *T. cruzi* populations within triatomine species in this region. Notably, the divergent TcI lineage detected in Florida coincides with the geographic distribution of *Triatoma ambigua*, a vector species recently revalidated as distinct from *T. sanguisuga*. This correspondence suggests that vector evolutionary history, alongside geographic isolation on the Florida peninsula, may contribute to shaping contemporary patterns of *T. cruzi* genetic diversity in the southeastern United States.

The multilocus approach using deep sequencing enabled the identification of mixed infections, mitochondrial introgression, and multiclonality among the analyzed specimens. While the presence of shared genetic variants across different populations reinforces the predominantly clonal nature of *T. cruzi*, the detection of mitochondrial introgression provides clear evidence of genetic exchange between DTUs; specifically, the transfer of mitochondrial genetic material from TcIV to TcI, an event inconsistent with strict clonality. Finally, our study underscores the value of a multilocus strategy targeting single-copy genes combined with high-throughput sequencing to provide a more comprehensive assessment of *T. cruzi* genetic diversity. This approach minimizes biases associated with multi-copy sequences and conventional methods that primarily detect dominant haplotypes. A more exhaustive characterization of *T. cruzi* infections is essential for improving our understanding of their ecological and epidemiological dynamics within hosts.

## Acknowledgments

We would like to acknowledge our collaborators who contributed to the sampling and field collection efforts: Robert L. Smith, Chanakya R. Bhosale, Carson W. Torhorst, and Walter Roachell. We are also grateful to the community scientists in Florida for their help identifying suitable regions for triatomine sampling.

## Data Availability

The raw reads have been deposited under BioProject (https://www.ncbi.nlm.nih.gov/bioproject/) accession number PRJNA1283157 and BioSample (https://www.ncbi.nlm.nih.gov/biosample) accession numbers: SAMN49683252 to SAMN49683320. The Unique genetic sequences (haplotypes) were deposited in GenBank (https://www.ncbi.nlm.nih.gov/genbank/) accession numbers: PX229625 to PX229674.

## Supporting information

**S1 Fig. *Trypanosoma cruzi* phylogeny based on 17c nuclear loci.** Branch values indicate the statistical support from the maximum likelihood analysis (first value) and the posterior probability from the Bayesian inference (second value). Sequences found in this study are highlighted in bold within the tree. Sequences followed by an asterisk (*) correspond to assemblies from hybrid DTU strains, while those with a black dot (●) at the end of the name correspond to sequences obtained by Sanger sequencing. Input data: 85 sequences with 440 nucleotide sites. Best-fit model according to BIC: K3P.

**S2 Fig. *Trypanosoma cruzi* phylogeny based on 17d nuclear loci.** Branch values indicate the statistical support from the maximum likelihood analysis (first value) and the posterior probability from the Bayesian inference (second value). Sequences found in this study are highlighted in bold within the tree. Sequences followed by an asterisk (*) correspond to assemblies from hybrid DTU strains, while those with a black dot (●) at the end of the name correspond to sequences obtained by Sanger sequencing. Input data: 111 sequences with 333 nucleotide sites. Best-fit model according to BIC: K3P.

**S3 Fig. *Trypanosoma cruzi* phylogeny based on 18c nuclear loci.** Branch values indicate the statistical support from the maximum likelihood analysis (first value) and the posterior probability from the Bayesian inference (second value). Sequences found in this study are highlighted in bold within the tree. Sequences followed by an asterisk (*) correspond to assemblies from hybrid DTU strains, while those with a black dot (●) at the end of the name correspond to sequences obtained by Sanger sequencing. Input data: 81 sequences with 412 nucleotide sites. Best-fit model according to BIC: K2P.

**S4 Fig. *Trypanosoma cruzi* phylogeny based on pyruvate dehydrogenase kinase (*TcPDK*) nuclear loci.** Branch values indicate the statistical support from the maximum likelihood analysis (first value) and the posterior probability from the Bayesian inference (second value). Sequences found in this study are highlighted in bold within the tree. Input data. Sequences followed by an asterisk (*) correspond to assemblies from hybrid DTU strains, while those with a black dot (●) at the end of the name correspond to sequences obtained by Sanger sequencing. Input data: 81 sequences with 414 nucleotide sites. Best-fit model according to BIC: HKY+F.

**S5 Fig. *Trypanosoma cruzi* phylogeny based on Pumilio/PUF RNA-binding protein 1 (*TcPUF1*/ RB) nuclear loci.** Branch values indicate the statistical support from the maximum likelihood analysis (first value) and the posterior probability from the Bayesian inference (second value). Sequences found in this study are highlighted in bold within the tree. Input data. Sequences followed by an asterisk (*) correspond to assemblies from hybrid DTU strains, while those with a black dot (●) at the end of the name correspond to sequences obtained by Sanger sequencing. Input data: 79 sequences with 370 nucleotide sites. Best-fit model according to BIC: K3P.

**S6 Fig. TCS haplotype networks for the five nuclear markers of *T. cruzi* in Triatominae species from the southern United States. (A) Haplotypes grouped by state. (B) Haplotypes grouped by species.**

**S1 Table. Whole-genome sequencing data for *Trypanosoma cruzi* strains used in this study for the genotyping and classification of *T. cruzi* Discrete Typing Units (DTUs).**

**S2 Table. Metadata for the *Trypanosoma cruzi*–positive triatomine specimens from natural populations in the southern United States included in this study.**

**S3 Table. Unique genetic sequences (haplotypes) identified for five nuclear loci among *T. cruzi* populations within Triatominae species across the southern United States.**

**S4 Table. Unique genetic sequences (haplotypes) identified for *cytb* among *T. cruzi* populations within Triatominae species across the southern United States.**

**S5 Table. Analysis of Molecular Variance (AMOVA) for *T. cruzi* of Triatominae species across the southern United States.**

